# DNA replication timing reveals genome-wide features of transcription and fragility

**DOI:** 10.1101/2024.11.25.625090

**Authors:** Francisco Berkemeier, Peter R. Cook, Michael A. Boemo

## Abstract

DNA replication in humans requires precise regulation to ensure accurate genome duplication and maintain genome integrity. A key indicator of this regulation is replication timing, which reflects the interplay between origin firing and fork dynamics. We present a high-resolution (1-kilobase) mathematical model that maps firing rate distributions to replication timing profiles across various cell lines, validated using Repli-seq data. The model effectively captures genome-wide replication patterns while identifying local discrepancies. Notably, regions where the model and data diverge often overlap with fragile sites and long genes, highlighting the influence of genomic architecture on replication dynamics. Conversely, regions of high concordance are associated with open chromatin and active promoters, where elevated firing rates facilitate timely fork progression and reduce replication stress. By establishing these correlations, our model provides a valuable framework for exploring the structural interplay between replication timing, transcription, and chromatin organisation, offering new insights into mechanisms underlying replication stress and its implications for genome stability and disease.

## Introduction

Accurate DNA replication is essential for faithfully duplicating genetic information, ensuring its preservation for future generations (Gefter, 1975). In humans, replication occurs during S phase when multiple discrete chromosomal sites, termed origins of replication (Leonard and Méchali, 2013), “fire” to initiate bidirectional replication forks—molecular machines that traverse the chromosome and replicate DNA (Waga and Stillman, 1998). These forks move in opposite directions, progressing until they encounter another fork, reach a chromosome end (Figure 1a), or face an obstacle (e.g., a bound protein or transcription complex; Mirkin and Mirkin (2007)). Intriguingly, each origin fires stochastically so firing sites and times differ from cell to cell. Despite this apparent randomness, consistent trends emerge so that different cell types have characteristic firing profiles (Rhind and Gilbert, 2013).

**Figure 1.**
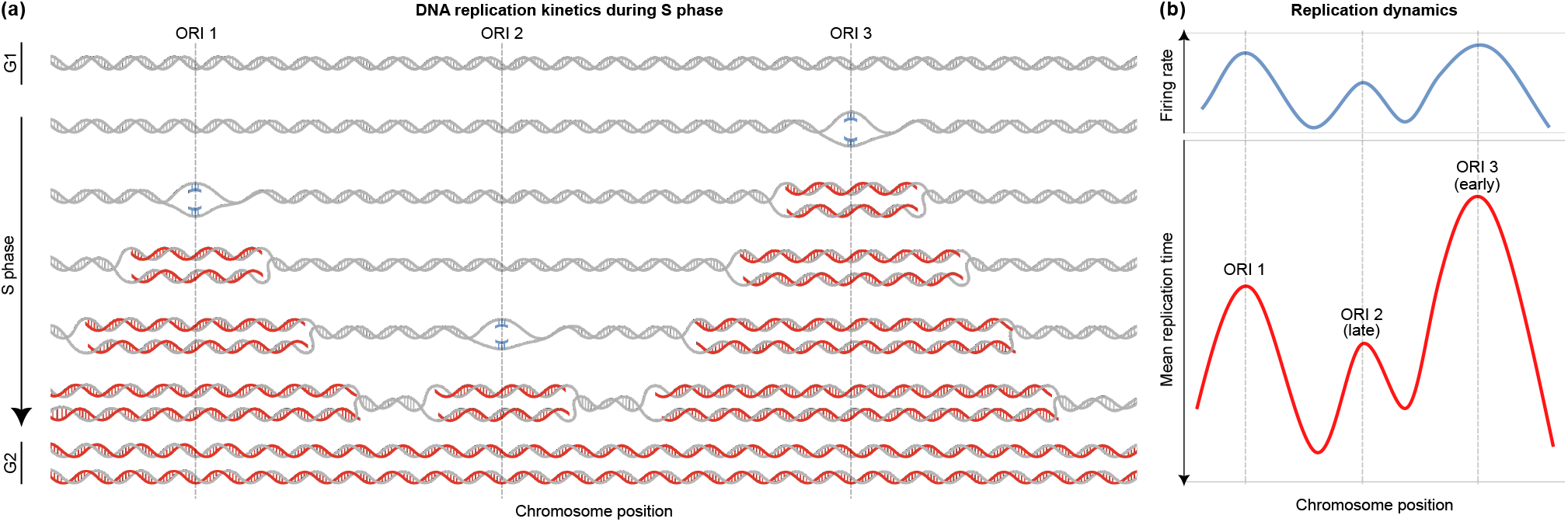
A kinetic model of DNA replication. **(a)** Replication initiates at specific origins that are fully licensed by the end of G1 phase. During S phase, replication forks progress bidirectionally from origins, passively replicating DNA until they merge with forks from adjacent origins or reach chromosome ends to complete replication and enter G2. In this example, three origins (ORIs 1, 2, and 3) fire at different times, with nascent DNA strands shown in red. At the end of replication, two identical copies of the original template are formed. **(b)** Illustration of the expected inverse but non-trivial correlation between firing rates (top) and replication timing (bottom, with an inverted y-axis). In a model where the firing time of each origin is an exponentially distributed random variable, the firing rate is the parameter of this distribution and tends to decrease as replication timing increases, indicating that regions with higher firing rates replicate earlier in S phase. Replication timing, measured by Repli-seq, shows the average replication time across a cell population, with peaks corresponding to potential origins. ORI 2 is in a late-replicating region, while ORI 3 replicates earlier, as indicated by their relative positions on the timing curve.

Replication timing is a critical marker of replication fidelity, reflecting the time it takes for a specific locus either to fire (if an origin) or to be passively replicated by an incoming fork. These timing profiles are closely associated with various chromatin structures (Marchal et al., 2019), as well as gene expression (Müller and Nieduszynski, 2017) and replication stresses (Briu et al., 2021). Furthermore, timing is linked to genetic variation (Koren et al., 2014) and cancer (where late or delayed replication often correlates with increased genomic instability; Woo and Li (2012)). Of particular interest are fragile sites, regions that are especially vulnerable to breakage due to replication stress, and are often found in late-replicating regions (Sinai and Kerem, 2018). These sites, and the long genes found within them, are often hotspots for the chromosomal rearrangements and deletions that arise in cancers and other genetic diseases (Smith et al., 2006).

Replication, transcription, and chromatin organisation are also intricately inter-connected, with each influencing the other (Sequeira-Mendes et al., 2009; Ehrenhofer-Murray, 2004; Turner and Woodworth, 2001). In particular, chromatin remodelling regulates the accessibility of regulatory factors, influencing both gene expression and replication. Open chromatin is strongly linked to transcriptional activity and plays a crucial role in replication timing (Guilbaud et al., 2011; Audit et al., 2009).

Although associations between genomic features are well-established, identifying site-specific or context-dependent differences remains a challenge. Experimental approaches often struggle to isolate individual variables, limiting our ability to disentangle the interplay between replication and other processes. To address these gaps, we develop a stochastic model that maps origin firing rates to replication timing, capturing variability across cell populations. By integrating data from RNA-seq (Marguerat and Bähler, 2010), ChIP-seq (Pepke et al., 2009), GRO-seq (Lopes et al., 2017), and a database of fragile sites (HumCFS; Kumar et al. (2019)), we provide a framework to explore how discrepancies between the model’s predictions and experimental data may reflect signatures of transcriptional activity, chromatin openness, and genomic fragility.

We begin our analysis with a fundamental inquiry: how accurately can a kinetic model of replication predict genome-wide timing? Our model acts as a null hypothesis, representing how replication should occur in the absence of perturbation from genomic features. The central aim is to identify loci where the model’s predictions diverge from experimental observations, highlighting regions that may experience replication stress or other anomalies. By deriving a closed formula for the expected time of replication at each genomic site, we establish a solid mathematical framework to support our computational simulations.

Our workflow is simple: using only timing data as input, along with minimal genomic parameters such as potential origin locations, the model determines firing rates and predicts timing profiles plus other key kinetic features like fork directionality and inter-origin distances. Researchers with replication timing data can use this model to rapidly generate precise replication dynamics profiles without extensive computational expertise, revealing factors that influence replication timing and genome instability across various contexts.

Despite significant advances in mathematical modelling (Jun et al., 2005; Jun and Bechhoefer, 2005; de Moura et al., 2010; Retkute et al., 2012), deriving a position-specific, data-fitted model that precisely links replication timing to origin firing has remained a challenge. While some approaches rely on neural networks to infer probabilistic landscapes of origin efficiency (Arbona et al., 2023), ours differs by deriving a closed-form relationship between timing and firing. Rather than relying on complex inference techniques, our model abstracts intrinsic firing rates without directly tying them to specific biological mechanisms such as licensing or activation. This allows a precise fit to observed timing data and enables simulation of genome-wide dynamics in a direct and interpretable manner. Our approach improves existing fitting methods by adopting a convolution-based interpretation of the timing programme. Using process algebras from concurrency theory (Boemo et al., 2020), we model replication forks and origins as interconnected entities, simulating their behaviour across the genome. The key contribution of this work is demonstrating how a theoretical description of replication timing helps uncover links between timing, genomic stability, and other essential genomic processes.

## Methods

### Modelling assumptions

We aim to identify and quantify genomic regions where replication timing deviates from model predictions, hereafter referred to as replication timing misfits, which may indicate potential sites of replication stress or instability. To accomplish this, we model the complex, nonlinear relationship between origin firing rates and replication timing (Figure 1b) and fit these rates to experimental timing data. This approach enables investigation using replication forks, origins, and DNA templates as the level of abstraction. In particular, we do not differentiate between leading and lagging strands, as the formation and joining of Okazaki fragments are not explicitly included in the model. By concentrating on the fundamental kinetics driving replication, we gain a clearer understanding of how origin firing influences outcomes.

Our model operates under several key assumptions. The time an origin fires is modelled as an exponentially distributed variable (independent of fork movement and firing of other origins), and fork movement as an exponentially distributed random variable (independent of origin firing and movement of other forks). We also assume a constant rate of fork movement throughout (no fork stalling at obstacles); then, forks advance smoothly until encountering another fork or chromosome end. This assumption avoids overfitting and indirectly emphasizes the role of origin firing.

The key variable is the origin firing rate. This encompasses origin licensing and activation, plus contributions of all other proteins and pathways within this process. While a strong assumption, it is justified by the fact that firing rates effectively capture the collective outcome of all these underlying processes without explicitly representing molecular detail. This makes the model both tractable and capable of producing accurate genome-wide predictions. We further sub-divide the genome into 1 kb intervals (sites), and assign to each a non-zero firing rate determined by a governing equation that links timing with firing. This resolution offers a balance between computational efficiency and biological realism. Although any site is a potential origin, passive replication and low firing rates ensure the expected sparsity of origins seen in the genome.

### Mathematical modelling of replication

Consider a DNA molecule with *n* discrete genomic loci, where each locus can potentially act as an origin that fires at rate *f* to initiate a fork that progresses bidirectionally with speed *v*, typically measured in kilobases per minute (kb/min). We aim to determine the average time required for a site to either initiate replication, or to be passively replicated by an approaching fork (i.e., its expected replication time). Initially, we assume that all origins fire at the same rate, *f*, but later relax this assumption to allow for variations in firing rates across different origins. In addition, by considering a sufficiently large chromosome, we ensure that effects of chromosomal ends are negligible. Nonetheless, the framework can easily be extended to account for such effects, though they are not critical for the broader analysis.

#### Expected time of replication

Let *T* be the time a site takes to fire or be passively replicated by a fork. We assume initially that all origins fire at the same rate, *f*. One may think of *T* as an explicit function of origin firing times *A*_*i*_, where 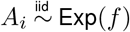. In particular, 𝔼 [*A*_*i*_] = 1*/f*. We index each site by its distance from the origin of interest, given by |*i*|. Notice that *i* = 0 corresponds to the focal origin, and *v* is interpreted as the number of replicated sites per time unit. We have

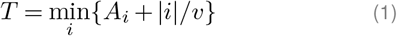

since it takes time |*i*|*/v* for a fork initiated at site *i* to reach the origin of interest. Next, we compute the cumulative distribution function. The minimum in Eq. (1) is greater than some *t* if all terms are, which occurs with probability

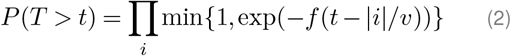

since *A*_*i*_ *>* 0 and 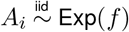. Hence, the expectation of replication time for any one site is given by

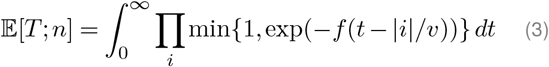

where the product is taken over all *n* sites. This integral can be partitioned across each interval for which |*i*| ≤ *vt* ≤ |*i* + 1|. Within these intervals, integrands adopt the form *ae*^−*bt*^, thereby permitting analytical evaluation. In the general case, the result depends on the parity of *n*. See Supplementary Note (SN) 1.1 for an explicit expression of 𝔼 [*T*; *n*].

As *n* → ∞, a general expression of the expected replication time for each origin can be written as

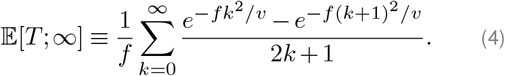

With *v* = 1.4 kb/min (Conti et al., 2007), Figure 2a shows the dynamics of 𝔼 [*T*; *n*] for increasing values of *n*. By relating Eq. (4) to the family of theta and Dawson functions (Tyurin, 2002; Temme, 2010), the following approximation holds (see SN1.2 for a detailed proof)

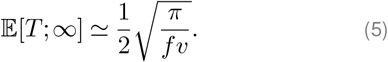

**Figure 2.**
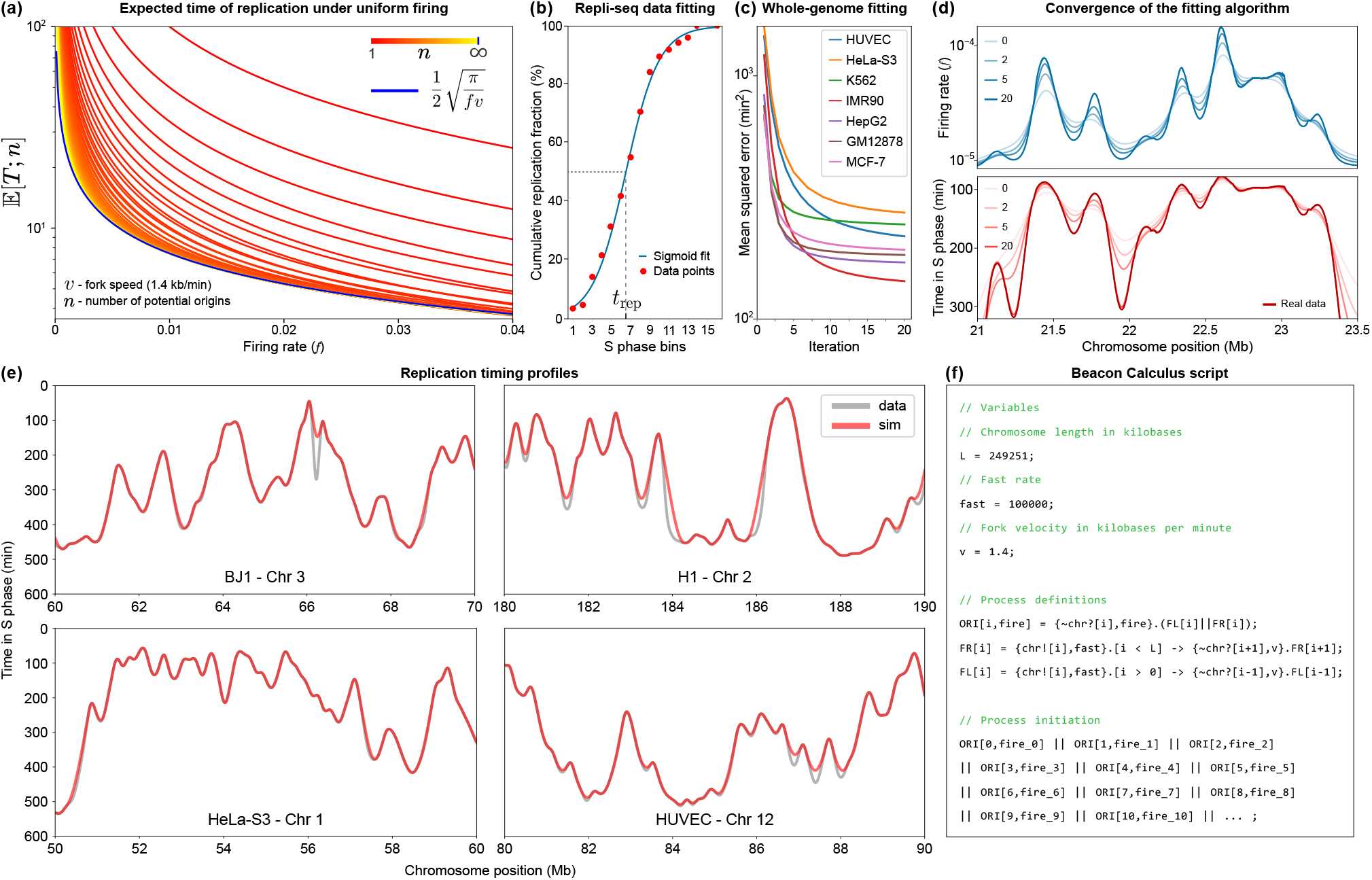
Fitting the model. **(a)** Replication asymptotics under uniform firing: logarithmic plot of the expected replication time, 𝔼 [*T*; *n*], as a function of the firing rate, *f*, and the number of potential origins, *n* (spaced at 1 kb intervals), for 1 ≤ *n <* ∞, with *v* = 1.4 kb/min. As *n* → ∞, 𝔼 [*T*; *n*] approximates an inverse power law (blue). **(b)** Curve fitting for cumulative replication in S phase. Red markers depict example data points from a high resolution Repli-seq heatmap that shows the cumulative percentage of completed replication across 16 S phase bins. The blue line is the curve fitted to this data, while the dashed grey line indicates the median replication time, *t*_rep_ (the instant in S phase when 50% of replication is achieved across the cell population). **(c)** Whole-genome mean squared error between simulated timing profiles and real data for 7 cell lines, in min2. Fitting each line took ∼ 3 minutes on a HPC platform (one CPU). **(d)** Progression of the fitting algorithm over 20 iterations for chromosome 2 in the BJ line on firing rates (above), with iteration 0 corresponding to the initial inverse power law estimate, given by Eq. (6), and the corresponding timing profile (below). **(e)** Observed (Repli-seq) timing against the simulated profiles for different lines and genomic regions. **(f)** Model written in the Beacon Calculus process algebra. Origin firing processes take their location, `i` (1-kb resolution), and firing rate `fire`, as parameters, triggering two replication fork processes, `FL` (left-moving) and `FR` (right-moving). Replication terminates when all locations have been replicated. The simulation begins by invoking the `ORI` processes, where `fire_i` corresponds to the firing rate values for each origin `i`, as determined by fitting Eq. (8).

Provided replication timing data {*T*_*j*_}_1≤*j*≤*n*_, we have the following inversion

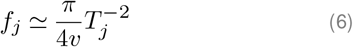

which provides a first estimate for the intrinsic firing rate of an origin, given its time of replication. Note that Eq. (6) is an approximation under the specific assumption that firing rates are uniformly constant across the genome, a simplification that, intriguingly, offers a reasonably accurate initial estimate for the firing rate distribution in most instances. The fidelity of this approximation is closely tied to fork speed *v* and the average of the timing dataset, topics that will be elaborated subsequently.

#### A generalisation

Experimental data support the idea that different origins fire at different rates (Rhind et al., 2010). While our introductory argument assumes a constant firing rate *f* across the genome, we should, in general, expect *A*_*i*_ ∼ Exp(*f*_*i*_). Then, the replication time definition in Eq. (1) should include the site-specific indexation, for 1 ≤ *j* ≤ *n*, as follows

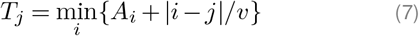

with indexes congruent modulo *n*, that is, |*i* −*j*|∈ ℤ*/nℤ* (see SN1). Following a similar argument, the general expression for 𝔼 [*T*_*j*_; ∞], with general firing rates {*f*_*i*_}, is approximately given by

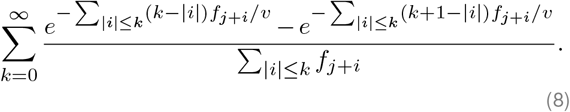

When *f*_*j*_ = *f*, ∀*j*, Eq. (8) is reduced to Eq. (4). While Eq. (8) holds true for an infinitely large genome, in practical terms this series can be limited to 0 ≤ *k* ≤ R < n/2, for some large enough *R*. This parameter represents the radius of replication influence: the distance within which neighbouring origins {*j* −*R*, …, *j* − 1, j + 1, …, *j* + *R*} are assumed to affect the timing of a focal origin *j*. In other words, while every firing origin does theoretically affect replication timing at any other location, this effect decays rapidly with distance from the origin of interest *j*. Numerically, the finite version of Eq. (8) should mimic the average replication timing obtained from computational simulations and it will be crucial in solving the fitting problem efficiently. Ideally, we would like to compute the rates {*f*_*j*_}_1≤*j*≤*n*_ as a function of the expectation of *T*_*j*_. Our goal is then to find a solution to Eq. (8), given data on *{𝔼* [*T*_*j*_; *n*]*}*, for large *n*. Alternative frameworks inspired by the analogy between DNA replication and crystal growth have been previously explored by Jun et al. (2005); Jun and Bechhoefer (2005); Jun and Rhind (2008), revealing other relevant replication metrics, such as inter-origin distances (Herrick et al., 2002). Our formulation extends these approaches by estimating origin firing rates from discrete replication timing data across the entire human genome, which is discussed next.

### Replication timing data

Replication timing data was sourced and processed from two key databases: Encyclopedia of DNA Elements (ENCODE; Hansen et al. (2010); Davis et al. (2018)) and high-resolution Repli-seq from Zhao et al. (2020). To ensure data consistency and reliability, extensive filtering and scaling steps were performed on all data sets. We analyse data from: HUVEC (human umbilical vein endothelial cells), HeLa-S3 (clonal derivative of the parent HeLa, an immortalised cervical cancer line), BJ (normal skin fibroblast), IMR90 (lung fibroblast), K562 (lymphoblast cells), GM12878 (lymphoblastoid line), HepG2 (hepatocellular carcinoma line), MCF-7 (breast cancer line), HCT (colorectal carcinoma line), plus H1 and H9 (embryonic stem cell lines). Data for HUVEC, HeLa, BJ, IMR90, K562, GM12878, HepG2, and MCF-7 cells were obtained from the ENCODE database using the GRCh37 (hg19) human genome assembly (Hansen et al., 2010; Davis et al., 2018), while data for HCT, H1, and H9 cells were sourced from high-resolution Repli-seq, using the GRCh38 (hg38) assembly (Zhao et al., 2020).

Regarding ENCODE Repli-seq, timing data from each cell line were analysed across 6 cell cycle fractions: G1/G1b, S1, S2, S3, S4, and G2, given as a wavelet-smoothed signal to generate a continuous portrayal of replication across the genome (Thurman et al., 2007). Importantly, we rescaled the original wavelet signal, initially normalised from 0 to 100, by a factor of 6 to better align with an approximately 8-hour S phase.

Following standard Repli-seq methods, we applied a sigmoidal fit to the cumulative replication fraction, *F*_rep_, to determine replication timing according to Zhao et al. (2020). We consider the median replication time, *t*_rep_, defined as the bin value *t* where *F*_rep_(*t*) = 50%, indicating that half of the cell population has completed replication (Figure 2b). Although Eq. (8) theoretically represents the mean replication timing, it aligns closely with the median observed in Repli-seq data, as replication timing distributions generally exhibit a near-symmetric sigmoidal pattern. Additionally, the median is more robust to experimental noise and outliers, making it a practical and reliable measure in high-throughput experiments. Although recent studies have determined telomere timing data (Massey and Koren, 2022b), we do not incorporate them into our analysis.

Repli-seq data shows consistent patterns across different cell lines. We present representative results from multiple lines (Figure 2e), but specific analyses may be more suitable for certain cases, depending on the availability and quality of the data. Although regions with repetitive sequences or low complexity are often mapped poorly using Repli-seq data (Hansen et al., 2010; Zhao et al., 2020), these regions account for ∼ 20% of the genome and show only a weak correlation with high-misfit regions (phi coefficient = 0.21). Therefore, we retain this data in our analysis, as its impact is minimal (see SN2.3).

### Fitting algorithm

We develop an algorithm to efficiently fit genome-wide replication timing data, processing over 3,200,000 potential origins per genome by leveraging the mean-field dynamics captured by Eq. (6). While every site is treated as a potential origin, the algorithm effectively suppresses many by assigning them negligible firing rates, reflecting the selective activation of origins in the genome. For large *n*, Eq. (8) provides an excellent estimate, allowing us to apply a fitting algorithm directly to the theoretical expectation, rather than relying on averaged outputs from multiple simulations. The radius of neighbouring influence, *R*, may be refined for optimisation. At each site *j*, we set

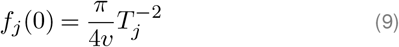

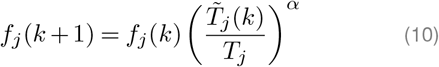

where 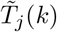 is the average replication time at iteration *k*, via Eq. (8), and *T*_*j*_ is the real data value. The parameter *α* is adjusted to guarantee convergence towards a stable firing rate distribution. The extent of each misfit is measured by the squared difference (fit error) between observed and expected replication timing, in min^2^, at each site. Numerical results and Eq. (6) suggest *α* = 2 is a reasonable compromise between error minimisation and speed. The optimised algorithm considers a convolution interpretation of Eq. (8) (SN2.2). Remarkably, the firing rates for each cell are fitted in an average time of approximately 3 minutes using 1 CPU on a high-performance computing platform equipped with Intel Ice Lake architecture (Figures 2c-e).

### Simulations

To simulate replication, we use Beacon Calculus (`bcs`), a process algebra designed for simulating biological systems (Boemo et al., 2020). Within this framework, replication is modelled using three core processes: replication origins (`ORI`), left-moving forks (`FL`), and right-moving forks (`FR`). Each process is associated with a specific position on the chromosome of length `L`, and origins have an additional parameter, the firing rate, `fire`, or *f* in our model (Figure 2f).

In `bcs`, *v* is understood as the rate of replication by a moving fork, which is held constant. This differs from the constant fork speed assumption underlying Eq. (8). Specifically, in the `bcs` case, the time *F*_*k*_ required for a fork to replicate *k* consecutive sites follows an Erlang(*k, v*) distribution, meaning that 𝔼 [*F*|*i*−*j*|] = |*i* −*j*|*/v*, which mirrors the approximation used in Eq. (7). Therefore, when averaged over a sufficiently large number of simulations, stochastic deviations in numerical simulations become negligible and they do not compromise the broader analysis or conclusions.

To track the progress of replication, the model marks regions of the chromosome that have been replicated, allowing us to monitor replication dynamics accurately. In all `bcs` simulations, fork speed was set to 1.4 kb/min (Conti et al., 2007), and results were averaged over 500 simulations, with the radius of influence set to *R* = 2,000 kb, as previously defined.

## Results

### Predicting genome-wide replication

After assigning the time of replication (determined using Repli-seq data) to every 1 kb segment of the genome in 11 different human cell lines, site-specific firing rates are fit to the data via Eq. (8). Then, replication is simulated using Beacon Calculus (`bcs`), a concise process algebra ideal for concurrent systems. Finally, we explore patterns of replication seen after averaging 500 simulations for each of the 11 lines (Figure 3a).

**Figure 3.**
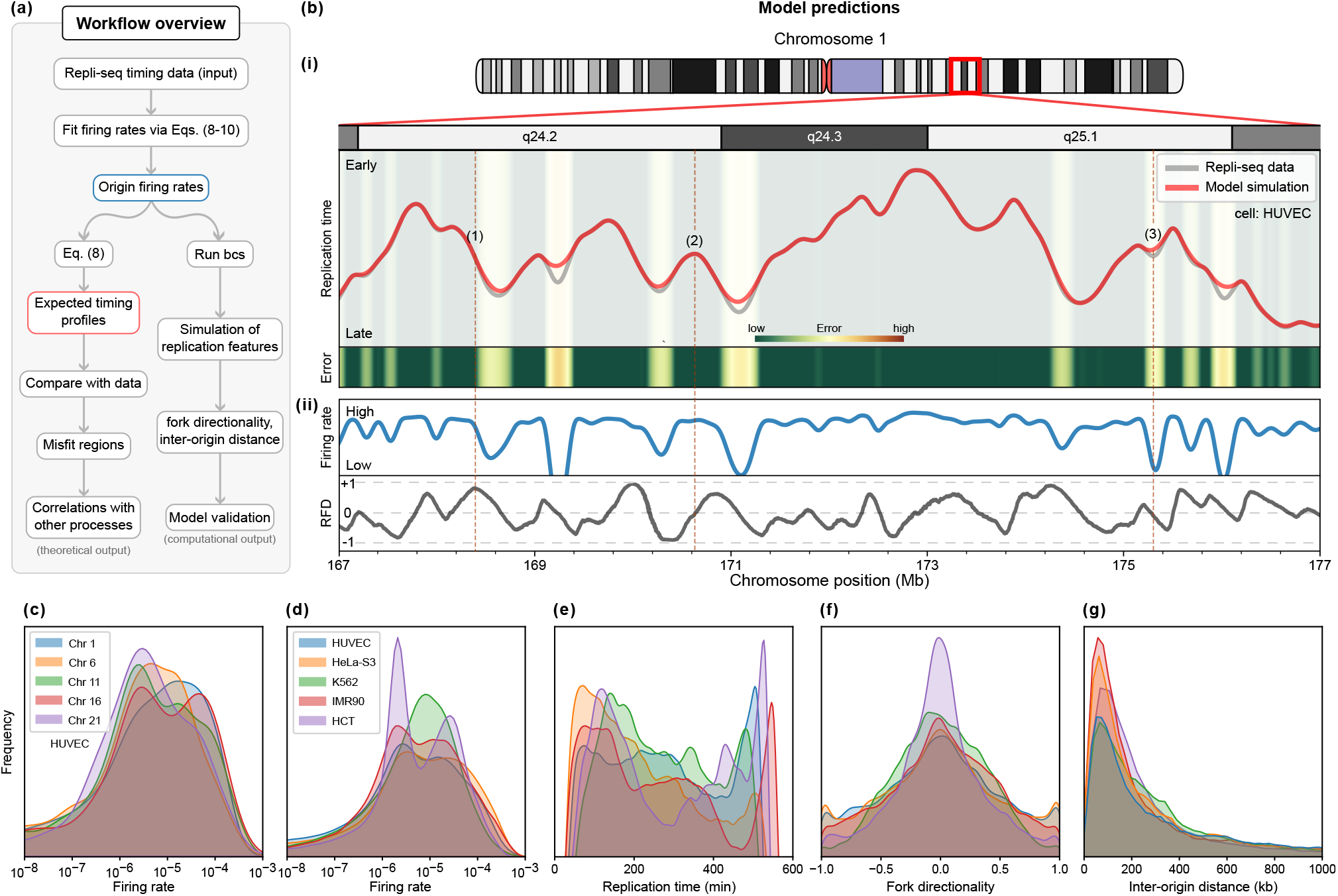
Predicting genome-wide features of replication. **(a)** Overview of the main model and analysis. Starting with Repli-seq timing data, origin firing rates are fitted through Eqs. (8-10). These rates generate expected timing profiles for comparison with experimental data to identify regions of timing misfits and fork stalling, which are analysed for correlations with other genomic processes. Simulations of replication features, such as fork directionality and inter-origin distances, validate the model against the literature. **(b)** Example of main modelling outputs from a region in HUVECs. **(i)** Replication timing of both experimental and simulated data, and the magnitude of the misfit (error) for replication timing in a region where replication forks often stall; this leads to elevated errors that the model struggles to capture accurately. **(ii)** Inferred origin firing rates, and fork directionality, scaled between -1 (leftward) and +1 (rightward). We highlight three regions of interest: (1) A passively replicated site predominantly replicated by rightward-moving forks (RFD ∼ 1); (2) A likely origin, characterised by a high firing rate and an RFD of 0; (3) A poorly fitted region between two origins with a low firing rate determined by the fitting algorithm with RFD of 0 (an equal likelihood of replication by leftward- and rightward-moving forks). **(c)** Kernel density estimate (KDE) of firing rate distributions across selected chromosomes in HUVECs. **(d-g)** KDEs comparing genome-wide features—including firing rates, replication timing, fork directionality, and inter-origin distances—across different cell lines. All distributions align with experimental observations. Areas under curves are equal to 1, while y-axis values are omitted to emphasize relative shapes and distributions rather than absolute magnitudes.

We begin by comparing experimental timing profiles to those obtained from Eq. (8). Note that this is equivalent to averaging the timing profiles from a large number of `bcs` simulations, which also allows us to save significant computational resources when computing timing alone. An example for chromosome 1 in HUVECs is shown in Figure 3b.i. As expected, some regions replicate early (e.g., around 173 Mb) and others late (e.g., around 171 Mb). There is generally excellent concordance between the model’s output compared with the experimental input. Our focus is on regions with high misfit error (shaded yellow and red areas; Figure 3b.i), where concordance breaks down because Eq. (8) predicts an earlier replication time. Instances where it predicts later replication compared to data are exceedingly rare, underscoring the algorithm’s reliance on higher firing rates to achieve the best fit with theoretical expectations.

While firing rates are directly inferred from Eq. (8), replication fork directionality (RFD) is calculated as the proportion of cell cycles (or `bcs` simulations) in which a given site is replicated by rightward versus leftward forks. RFD values range from -1 (always replicated by leftward forks) to +1 (always replicated by rightward forks), with intermediate values indicating a mix of replication directions across simulations (Figure 3b.ii).

To validate the model, we examine global distributions of multiple features. Despite little variation in firing in HUVECs (Figure 3c), HCT exhibits a pronounced bimodal pattern, likely driven by differences in data sources (Figures 3d-e; Zhao et al. (2020)), which may affect how replication timing and origin firing rates are captured. Regarding RFD, our results demonstrate a balanced bidirectional fork movement, with fork directionality symmetrically distributed and accumulating around zero, indicating efficient replication progression (Figure 3f). This pattern aligns with recent quantifications of fork directionality in human cells (Anderson et al., 2024). While determining inter-origin distances (IOD) is straightforward from our simulations, doing so from DNA-fiber experiments remains challenging due to technical limitations and potential biases (Quinet et al., 2017; Técher et al., 2013). Nevertheless, simulations show a concentration of IODs within the commonly observed range of 100–200 kb (Figure 3g; Conti et al. (2007)).

Although these results validate the model against established metrics, its broader ability to simulate other features, like replicon lengths and active fork numbers, highlights its value in capturing the full spectrum of replication dynamics. The most compelling insight, however, comes from examining regions where the model’s predictions diverge from data, as these discrepancies may coincide with critical sites of genomic instability, revealing areas of unique biological interest, which we address next.

### Hotspots of instability

We now determine genome-wide error profiles in all 11 cell types (Figure 4a illustrates those for chromosome 1). Remarkably, some of the regions that fit poorly are found in all cell lines (despite using different genome builds); this underscores the robustness of profiles across cell types (Bracci et al., 2023; Müller and Nieduszynski, 2012). Replication timing and firing rates are strongly negatively correlated (Spearman’s rank correlation of ∼ -0.89; Figure 4b); regions with higher firing rates tend to replicate earlier. Late-replicating regions also have a wide spread of low firing rates, reflecting a pattern captured by the fitting algorithm. Additionally, the lowest errors are seen in the earliest replicating regions, moderate ones in both early- and late-replicating regions, and the highest are distributed throughout mid-to-late S phase (Figure 4c). This suggests misfits increase as S phase progresses and fewer firing events occur. Low firing rates are also associated with high errors (Figure 4d; note the branched profile, reflecting difficulties in accurately modelling high-to-low firing rate transitions). Timing misfits are predominantly concentrated in late-replicating regions (Figure 4e). This is consistent with prior results suggesting that the replication machinery encounters more obstacles towards the end of S phase (Branzei and Foiani, 2010; Colicino-Murbach et al., 2024). Additionally, errors exceeding 10^4^ (min^2^) are more frequent in non-coding regions compared to coding ones, indicating a potential vulnerability of non-coding DNA to replication stress. Misfits also vary between cell lines, with HCT displaying a distinct pattern likely due to differences in data processing (Figure 4f; see Methods). Similar disparities were observed previously (Figure 3d), hinting at the potential for cell line-specific analyses to offer further insights. However, given our focus here, we leave a detailed analysis of these dynamics for future exploration.

**Figure 4.**
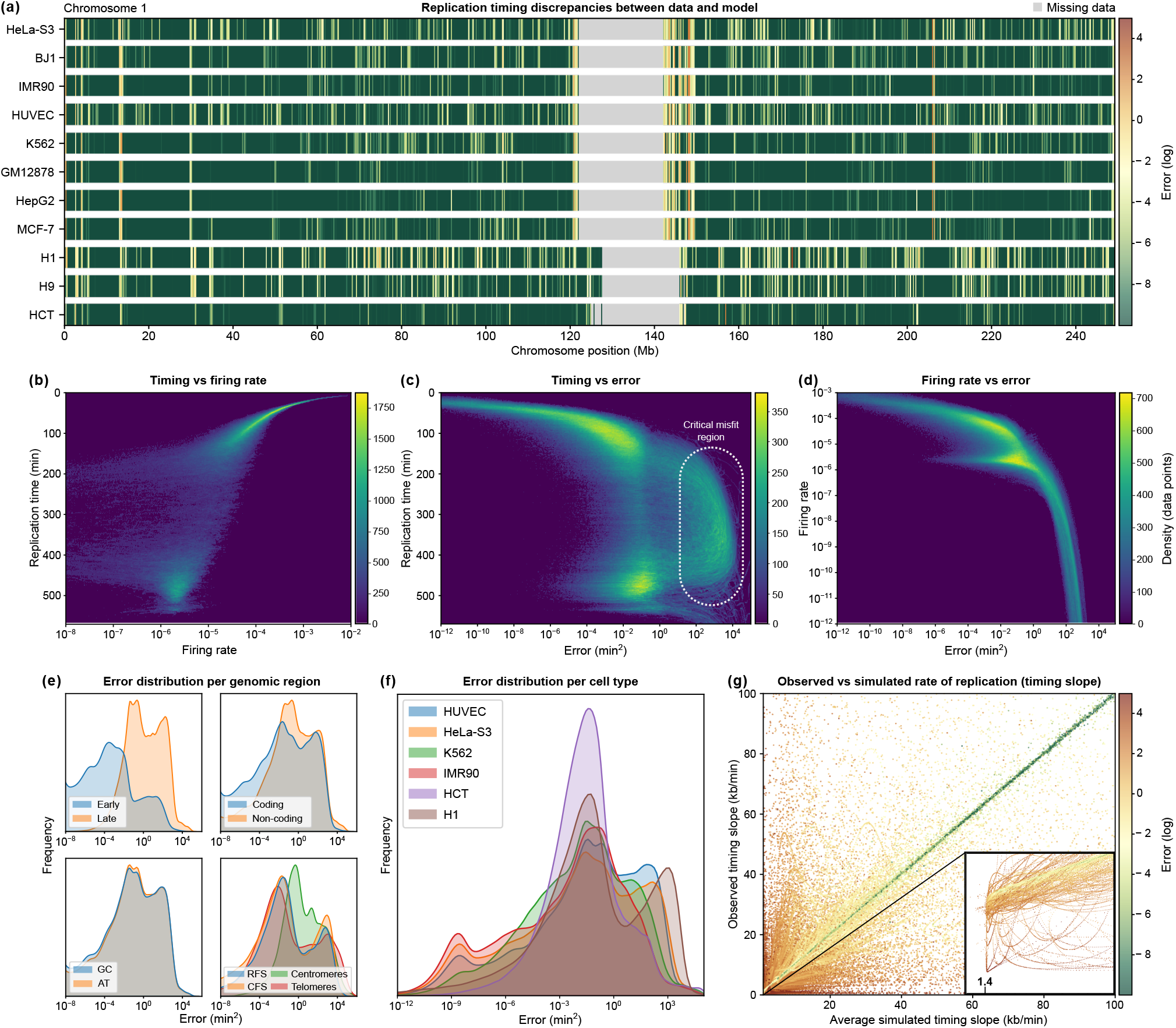
Detecting discrepancies in replication timing determined experimentally and in simulations. **(a)** Normalised error plots (red - high error, green - low error) highlighting deviations between simulated and experimental replication timings (chromosome 1 in various human cell lines). Grey areas: missing or unavailable data. **(b-d)** Density scatter plots illustrating key relationships in H1 cells (averages of 500 simulations). We look at the pairwise combinations of three variables: replication time, firing rate, and error. Density is represented as the number of data points per pixel on an 80 DPI (dots per inch) resolution across all plots. In **(b)**, the inverse correlation between replication timing and firing rate is evident, with greater variability in firing rates late in S phase. **(c)** highlights the relationship between replication timing and error, showing that high errors are distributed throughout S phase (dotted oval). **(d)** illustrates the branching relationship between firing rate and error. **(e)** Error distributions in HUVECs. Early-replicating regions tend to be error prone, coding and non-coding regions have broadly similar patterns, AT-rich regions are much like GC-rich ones, and different genomic regions have characteristic profiles. **(f)** Genome-wide error profiles in different cells. **(g)** Scatter plot comparing average simulated timing slope, indicative of the progress of replication over time, against observed data, color-coded by associated error. The zoomed-in region at [1.2, 2] *×* [0, 2] kb/min highlights the 1.4 kb/min lower bound on the simulated slope. Each dot represents a simulated-observed data pair, with the strand-like continuity arising from the high resolution of our 1 kb model, where proximity between adjacent pairs reflects the minimal positional shifts captured at this scale.

In regions with infrequent origin firing, the slope of the timing curve—representing the rate of replication changes over time—is primarily governed by fork speed, establishing an effective lower bound of 1.4 kb/min (Figure 4g). This constraint becomes most evident in regions where observed slopes fall below such a bound, resulting in error accumulation around slower-replicating areas. Origin competition, where nearby origins fire at similar times, further compounds these errors, producing timing ‘valleys’ between origin firing peaks. These patterns highlight regions of potential stress, suggesting areas for further study.

### Fragile sites and long genes

Fragile sites are cytogenetically defined gaps and breaks in metaphase chromosomes (Li and Wu, 2020); examples include FRA3B (Letessier et al., 2011) and FRA16D (Palakodeti et al., 2004). They are often seen after partially inhibiting DNA synthesis or applying other replicational stresses (Glover et al., 2017). They also contain few origins (Sinai and Kerem, 2018), and probably arise due to fork stalling or collapse (Kaushal and Freudenreich, 2019). Fragile sites can be broadly categorised into common fragile sites (CFSs) present in the whole population and rarer ones (RFSs) found in only a few individuals. We consider both classes, using locations from the HumCFS database (Kumar et al., 2019) and gene locations from the GENCODE Genes track v46 (Frankish et al., 2023).

As seen in Figure 4c, replication timing misfits are most pronounced in mid-to-late S phase, where our model struggles to capture timings accurately. Regions such as centromeres and telomeres, as well as most fragile sites often map to regions of high error, particularly during late S phase (Figures 5a-d). FRA3B and FRA16D show even higher median misfit lengths, suggesting these regions are especially challenging for the model to fit (Figure 5e). Similarly, long genes in fragile sites, such as *CNTNAP2, LRP1B*, and *FHIT*, also exhibit substantial error (Figures 5f-g). Under a certain error threshold, long genes overlapping with misfit regions, including those in fragile sites, may be easily identified through our model (Figure 5h). Approximately 30% of these genes overlap fragile sites, reinforcing the established link between late replication timing and the demands of transcribing long genes. Notably, chromosomes 15, 20, and 22 show no significant misfits associated with fragile sites, likely due to a lower abundance or reduced activity of fragile sites on these chromosomes. While not all fragile sites follow this pattern, the observed timing misfits at long genes suggest broader genomic regulation factors that may underlie replication stress. Fragile sites are cell type-specific (Letessier et al., 2011; Le Tallec et al., 2011), yet we observed consistent trends in fragility-misfit correlations across all 11 lines analysed, with H1 cells used as an illustrative example; this consistency suggests conserved replication dynamics, with the implications of cell type-specific variation addressed in the Discussion.

**Figure 5.**
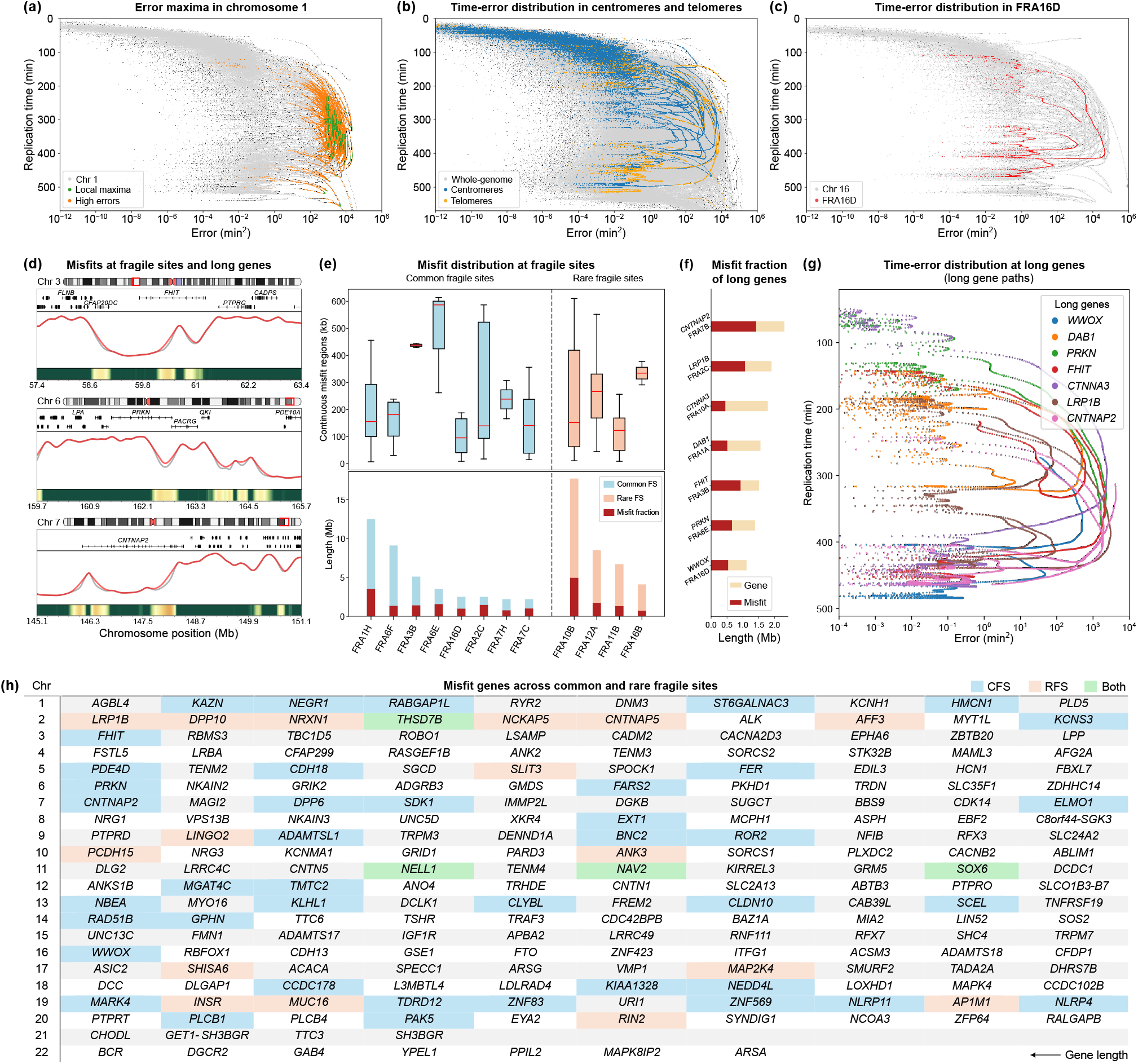
Timing errors in fragile sites and long genes. **(a)** Replication timing vs. error on chromosome 1 in H1, highlighting regions with local maxima in error and neighbouring high-error zones (within a 300 kb radius). The threshold for identifying local maxima in errors is set at 10^2.8^ (min^2^). Each dot represents an error-timing data pair, with the strand-like continuity arising from the high-resolution of our 1 kb model. **(b)** Genome-wide scatter plot displaying replication timing vs. error, with specific focus on centromeres (blue) and telomeres (orange), alongside the whole-genome data. **(c)** Scatter plot for chromosome 16, zooming in on FRA16D, revealing a continuous error path in mid-to-late replication, near the *WWOX* gene. **(d)** Examples of misfit regions detected by the model across three different chromosomes (3, 6, and 7). Each panel shows the chromosome ideogram, gene locations, and a comparison between the observed data (grey) and model predictions (red), as well the associated error. Notably, misfit regions overlap with long genes such as *FHIT* (Chr 3), *PRKN* (Chr 6), and *CNTNAP2* (Chr 7). **(e)** Misfit distribution for common (blue) and rare (pink) fragile sites. Top: length (in Mb) of continuous misfit regions. Bottom: misfit fraction relative to site length. **(f)** Misfit fraction analysis of the largest genes within fragile sites. **(g)** Scatter plot of replication timing vs. error trajectories for long genes, highlighting error accumulations based on gene size and location within fragile sites. **(h)** Table showing the 10 longest genes misfitted by the model across all chromosomes, ranked from largest to smallest (left to right). Genes intersecting fragile sites are highlighted in different colors: blue - common fragile sites, pink - rare fragile sites, green - both. All plots refer to H1 with Repli-seq reads aligned to hg38.

### Transcription and chromatin state

Transcription and replication have long been recognised to interact in complex and sometimes conflicting ways, particularly at fragile sites (Knott et al., 2009). Previous studies show that transcription-dependent barriers can obstruct fork progression, leading to stalling or collapse, while long genes associated with CFSs often initiate poorly, forcing forks to traverse longer distances from adjacent origins to delay replication completion (Blin et al., 2019). This delay is particularly pronounced in transcriptionally active regions. However, this is not always the case, as chromatin structure can play a more dominant role in timing discrepancies. Building on the previous results, we now turn our attention to interactions between transcription, chromatin structure, and replication.

Regulatory elements like active promoters and enhancers are marked by histone modifications such as H3K4me3 (Huang et al., 2019), DNase I hypersensitivity (DHS), and transcription-factor binding, detected using ChIP-seq (Cockerill, 2011; Young et al., 2011). By integrating data from ChIP-seq, RNA-seq, and GRO-seq (Cockerill, 2011; Crawford et al., 2004; Marguerat and Bähler, 2010; Lopes et al., 2017), we assess how these markers are associated with replication timing.

Regions with high GRO-seq signals align with peaks in H3K4me3 and DHS signals; they exhibit lower timing errors and higher firing rates (Figure 6a). Spearman rank correlation analyses reveal varying degrees of association between variables (Figure 6b). This method was chosen due to its suitability for non-normally distributed data and its ability to capture monotonic relationships, reflecting the ranked nature of our genomic features. Pearson and Kendall’s Tau tests were conducted for comparison (SN2.4). The consistently higher Spearman rank correlations indicate a strong monotonic relationship, particularly between DHS sites and firing rates, as well as between promoters and firing rates, revealing how chromatin accessibility facilitates replication initiation, even amid non-linear interactions.

**Figure 6.**
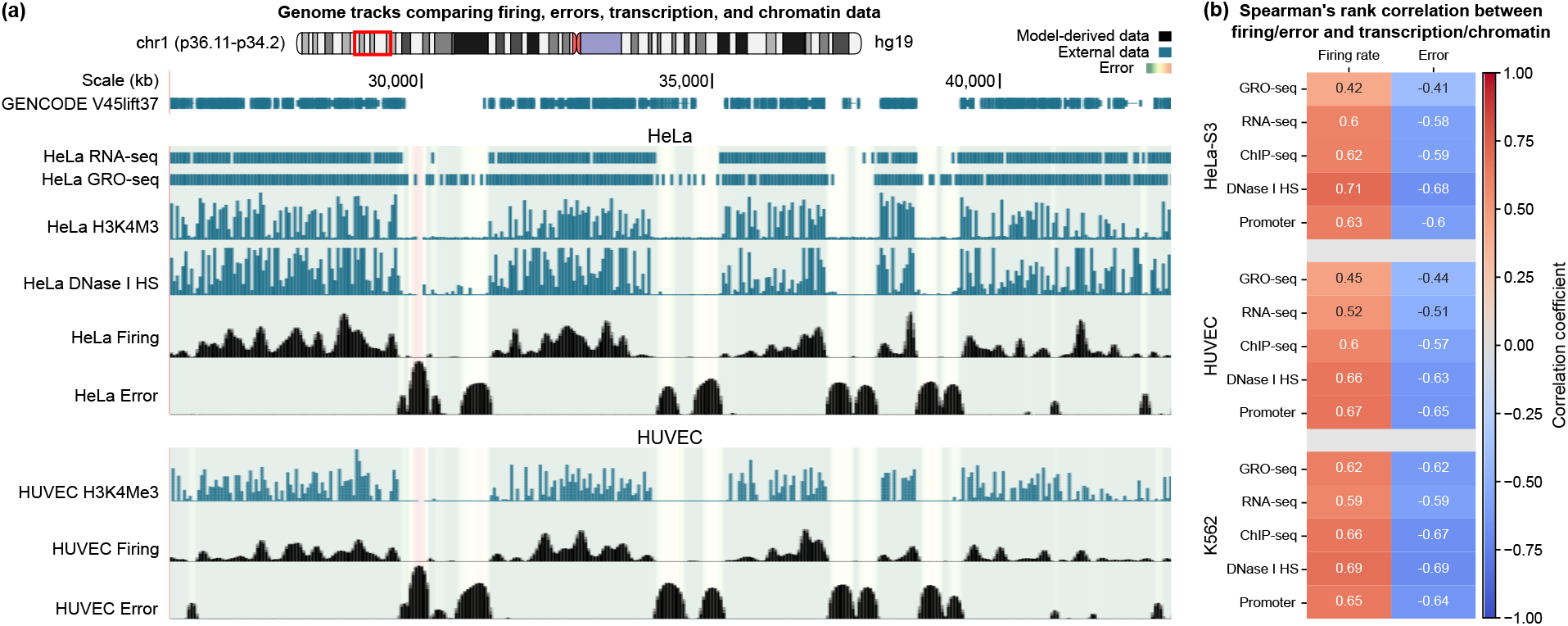
Replication timing discrepancies and firing rate profiles correlate with transcriptional and chromatin data. **(a)** Snapshot from the UCSC Genome Browser showing a detailed view of chromosome 1 (p36.11-p34.2) in HUVEC and HeLa (hg19). Various tracks compare transcriptional and chromatin data to misfit magnitude (error) and firing rate profiles obtained from our model (log-scale). Tracks include RNA-seq (marking mature mRNA levels), GRO-seq (nascent RNA), ChIP-seq for H3K4Me3 (promoters), and DNase I hypersensitivity (open chromatin). The error for each line is represented as a translucent heat map across tracks, with colours ranging from green (good fit) to yellow/red (poor fit). **(b)** Heatmap displaying the Spearman correlation coefficients between origin firing rates and fit errors with transcriptional and chromatin features for HeLa, HUVEC, and K562. All tests returned p-value *<* 10^−15^.

We observed a moderate to strong negative correlation between GRO-seq and misfits across all lines. This suggests that the replication machinery may encounter fewer impediments in regions with active transcription. A possible explanation is that transcriptionally active regions are more likely to be in an open state, reducing mechanical barriers to fork progression and lowering the chances of replication stress. Moreover, transcription factor-binding sites have been shown to enhance DNA replication, as evidenced by studies demonstrating that these sites significantly increase replication efficiency (Turner and Woodworth, 2001).

Furthermore, origin density strongly correlates with promoter density (Sequeira-Mendes et al., 2009). This coevolution of replication and transcription regulatory regions further supports the idea that transcriptional activity not only facilitates replication but also influences the efficiency and organisation of origins in mammalian cells. The strong correlation between high origin firing rates and regions of active transcription, open chromatin, and promoters provides further insight into genome-wide coordination of replication and transcription.

Notably, putative origins are often located in open and early-replicating chromatin (Audit et al., 2009; Chen et al., 2019) that is well fitted by our model. This synchronisation between replication and transcription may prevent replication stress, particularly during late S phase, aligning with the observation that transcriptionally active euchromatin tends to replicate early, and silent heterochromatin late (Gilbert, 2002). Under replication stress, this coupling is adjusted, with initiation and termination sites shifting to maintain the balance between replication and transcription, highlighting the intricate coordination that sustains genome integrity.

## Discussion

In genome-wide simulations, our model effectively captured key replication dynamics, including replication timing, fork directionality, and inter-origin distances. Replication timing was fitted with high precision across most of the genome, with only a few regions where observations clearly deviate from simulations (Figure 2). While misfit distributions varied across different chromosomes and cell lines (Figure 3), late-replicating regions consistently exhibited higher misfit rates (Figure 4). This matches previous findings suggesting these regions are more prone to replication challenges. Firing rates were also strongly negatively correlated with timing misfits; regions with infrequent origin firing are more susceptible to timing deviations. Additionally, non-coding regions had a higher frequency of misfits, highlighting their potential vulnerability. Misfits were particularly enriched in fragile sites and long genes (Figure 5). Our analysis pools data from multiple studies and cell types (Kumar et al., 2019). While this provides a broad overview, it does not account for the cell type-specific replication programs that underlie fragile site expression. Fragile sites are influenced by transcriptional activity and replication timing, both of which vary between cell types (Le Tallec et al., 2011; Brison et al., 2019). For instance, fibroblasts and lymphoblastoid cells exhibit distinct replication initiation patterns, which affect the timing and extent of fragility (Letessier et al., 2011). We observed consistent trends in the correlation between fragility and replication timing misfits across all 11 lines analysed, with H1 cells used as an illustrative example. This consistency highlights the robustness of the approach in identifying conserved replication dynamics and suggesting candidate regions and genes of interest. Researchers with access to cell type-matched Repli-seq and fragility data could refine this framework to achieve more specific insights. The investigation of early-replicating fragile sites (ERFS), which are associated with highly transcribed genes (Barlow et al., 2013), will be an important topic for future work.

Although the model does not incorporate detailed molecular mechanisms, regions with high origin firing rates were nonetheless strongly associated with active transcription, open chromatin, and promoter activity (Figure 6). These findings align with established knowledge, validating the model and underscoring its robustness. Notably, many misfit regions overlap with known fragile sites or distinct genomic locations, leading to the hypothesis that the model can refine the definition of fragile sites, distinguishing smaller, more nuanced regions of fragility, or even identifying novel sites prone to replication stress. Such predictions highlight the model’s utility in uncovering unexplored genomic vulnerabilities, warranting further experimental validation.

Our approach has various limitations. For instance, we simplistically assume that each origin fires independently of others, which may not capture the full complexity of origin licensing and activation. However, this simplification allows the model to fit human Repli-seq data rapidly, making it a practical tool for genome-wide analyses. Even so, in reality, a multiplicity of factors (e.g., ORC, Cdc6, and MCM proteins) regulate complex pathways of origin licensing, while later checkpoints and stress response pathways influence cell-cycle progression (Boos and Ferreira, 2019). Another limitation is that we take no account of higher-order genome structure, but could incorporate data from, for example, Hi-C (Gindin et al., 2014) and the position of R-loops, hairpins and G-quadruplexes that are known to obstruct replication and cause transcriptional conflicts (García-Muse and Aguilera, 2016). Furthermore, our model could highlight the relationship between origins and DNA break clusters, such as those found at timing transition regions, which are prone to replication-transcription conflicts and genome instability (Corazzi et al., 2024). A third limitation is that adapting the model for species like *Saccharomyces cerevisiae*, that have small genomes with precisely located origins, would necessitate adjusting assumptions. A feasible generalisation to achieve this involves setting low firing rates at specific locations and modifying the radius of influence in Eq. (8) to account for the distance to chromosome ends. Additionally, Repli-seq data averages timing across populations, masking single-cell heterogeneity (Massey and Koren, 2022a; Dileep and Gilbert, 2018); future studies using single-cell data could reveal variability in replication dynamics.

An exciting application of the model involves exploring the impact of chemotherapies on replication dynamics, particularly those therapies that target the Replication Stress Response (RSR) pathway and its key signalling proteins (Berti and Vindigni, 2016). By simulating the inhibition of these proteins, the model could provide valuable insights into how these disruptions affect replication timing, origin firing, and potential cell death (Manic et al., 2018). This could facilitate prediction of which combination chemotherapies might provide cost-effective approaches to optimise cancer treatments.

## Data and code availability

The GRO-seq data used in this study were obtained from the Gene Expression Omnibus (http://www.ncbi.nlm.nih.gov/geo/) under the following accession numbers: GSE62046, GSE94872, and GSE60454. The source code for the main fitting algorithm, along with the replication timing fit error and origin firing rate bedgraph files, are hosted on the following GitHub repository: https://github.com/fberkemeier/DNA_replication_model.git. The Beacon Calculus simulations were performed using version 1.1.0 of `bcs`, available at https://github.com/MBoemo/bc (Boemo et al., 2020). Additional examples of `bcs` scripts and optimisation algorithms can be found in SN2. For any comments, suggestions, or questions, please contact.

## ACKNOWLEDGEMENTS

We thank Prof. Sarah McClelland (Barts Cancer Institute, Queen Mary University of London) and Dr. Mathew Jones (Frazer Institute, University of Queensland) for their constructive and insightful feedback, which significantly improved the manuscript. We also thank all members of the Boemo lab for their helpful discussions and comments. This work was made possible by the Leverhulme Trust Research Project Grant RPG-2022-028. It was performed using resources provided by the Cambridge Service for Data Driven Discovery (CSD3), operated by the University of Cambridge Research Computing Service (https://www.csd3.cam.ac.uk), and supported by Dell EMC and Intel through Tier-2 funding from the Engineering and Physical Sciences Research Council (capital grant EP/T022159/1) and DiRAC funding from the Science and Technology Facilities Research Council (https://dirac.ac.uk). Additional funding and career support was provided by a Rokos Postdoctoral Associate position at Queens’ College Cambridge to Francisco Berkemeier and a fellowship at St John’s College Cambridge to Michael A. Boemo.

