## Supplementary Information for "DNA replication timing reveals genome-wide features of transcription and fragility"

### Supplementary Note 1: Mathematical notes

##### 1.1. Expected time of replication

Without loss of generality, we assume a ring network (periodic DNA) to enforce symmetry of replication with respect to a focal origin. In a large genome, this periodic assumption has minimal influence across most regions, apart from the chromosome ends.

Let $T$ be the time a site takes to either fire (if it is a replication origin) or be replicated by an incoming fork. We can think of $T$ as an explicit function of the origin firing times $A_i$, where $A_i \overset{\text{iid}}{\sim} \text{Exp}(f)$. In particular, $\mathbb{E}[A_i] = 1/f$. We index each site by its distance from the origin of interest, given by $|i|$. Notice that $i = 0$ corresponds to the focal origin, and $v$ is interpreted as the number of replicated sites per time unit. We have

$$T = \min_i \{A_i + |i|/v\} \tag{S1}$$

since it takes time $|i|/v$ for a replication fork initiated at site $i$ to reach the origin of interest. Then,

$$P(T > t) = \prod_i P(A_i > t - |i|/v) = \prod_i \min\{1, \exp(-f(t - |i|/v))\} \tag{S2}$$

since $A_i > 0$ and $A_i \overset{\text{iid}}{\sim} \text{Exp}(f)$. Hence, the expectation of the replication time for any one site is given by

$$\mathbb{E}[T] = \int_0^\infty P(T > t)\, dt = \int_0^\infty \prod_i \min\{1, \exp(-f(t - |i|/v))\}\, dt. \tag{S3}$$

This integral can be partitioned across each interval for which $|i| \leq vt \leq |i+1|$. Within these intervals, the integrands adopt the form $ae^{-bt}$, thereby permitting analytical evaluation. A few particular cases include:

- One origin ($n = 1$):

$$\mathbb{E}[T;1] = \int_0^\infty e^{-ft} dt = \frac{1}{f} \tag{S4}$$

- Two origins ($n = 2$):

$$\mathbb{E}[T;2] = \int_0^{\frac{1}{v}} e^{-ft} dt + \int_{\frac{1}{v}}^\infty e^{-f(2t-1/v)} dt = \frac{1}{f}\left(1 - \frac{1}{2} e^{-\frac{f}{v}}\right) \tag{S5}$$

- Three origins ($n = 3$):

$$\mathbb{E}[T;3] = \int_0^{\frac{1}{v}} e^{-ft} dt + \int_{\frac{1}{v}}^\infty e^{-f(3t-2/v)} dt = \frac{1}{f}\left(1 - \frac{2}{3} e^{-\frac{f}{v}}\right) \tag{S6}$$

- Four origins ($n = 4$):

$$\mathbb{E}[T;4] = \int_0^{\frac{1}{v}} e^{-ft} dt + \int_{\frac{1}{v}}^{\frac{2}{v}} e^{-f(3t-2/v)} dt + \int_{\frac{2}{v}}^\infty e^{-f(4t-4/v)} dt = \frac{1}{f}\left(1 - \frac{1}{12} e^{-4\frac{f}{v}} - \frac{2}{3} e^{-\frac{f}{v}}\right) \tag{S7}$$

where $\mathbb{E}[T;n] \equiv \mathbb{E}[T]$ for each $n$. In the general case, the result depends on the parity of $n$. When $n$ is odd, for each $k$, there are $2$ origins at a distance of $k = 1, 2, \ldots, (n-1)/2$ from the origin of interest. Adding up these distances leads to

$$\mathbb{E}[T; n_{\text{odd}}] = \sum_{k=0}^{(n-3)/2} \int_{k/v}^{(k+1)/v} e^{-f((2k+1)t - k(k+1)/v)}\, dt + \int_{(n-1)/(2v)}^\infty e^{-f(nt-(n-1)(n+1)/(4v))}\, dt, \tag{S8}$$

where the last term is just the $k = (n-1)/2$ term of the sum with the upper limit replaced by $\infty$. Solving the integrals yields

$$\mathbb{E}[T; n_{\text{odd}}] = \frac{1}{f}\left[\sum_{k=0}^{(n-3)/2} \frac{e^{-fk^2/v} - e^{-f(k+1)^2/v}}{2k+1} + \frac{e^{-f(n-1)^2/(4v)}}{n}\right]. \tag{S9}$$

When $n$ is even, for each $k$ there are 2 origins at a distance of $k = 1, 2, \dots, (n-2)/2$, and then there is 1 origin at a distance of $n/2$. Again, we add up the distances, each twice, but since there is only one origin at a distance of $n/2$, the very last distance sum is $n^2/4$. So, we get

$$\mathbb{E}[T; n_{\text{even}}] = \sum_{k=0}^{(n-2)/2} \int_{k/v}^{(k+1)/v} e^{-f((2k+1)t - k(k+1)/v)} dt + \int_{n/(2v)}^{\infty} e^{-f(nt - n^2/(4v))} dt. \tag{S10}$$

Solving the integrals yields

$$\mathbb{E}[T; n_{\text{even}}] = \frac{1}{f} \left[ \sum_{k=0}^{(n-2)/2} \frac{e^{-fk^2/v} - e^{-f(k+1)^2/v}}{2k+1} + \frac{e^{(-fn^2/(4v))}}{n} \right]. \tag{S11}$$

Using the ceiling function $\lceil \cdot \rceil$ to handle parity, a general expression for each origin, and any $n$, is

$$\mathbb{E}[T; n] \equiv \frac{1}{f} \left[ \sum_{k=0}^{\lceil (n-3)/2 \rceil} \frac{e^{-fk^2/v} - e^{-f(k+1)^2/v}}{2k+1} + \frac{e^{-f(\lceil (n-1)/2 \rceil)^2/v}}{n} \right]. \tag{S12}$$

In particular,

$$\mathbb{E}[T; \infty] \equiv \lim_{n \to \infty} \mathbb{E}[T; n] = \frac{1}{f} \sum_{k=0}^{\infty} \frac{e^{-fk^2/v} - e^{-f(k+1)^2/v}}{2k+1} \tag{S13}$$

which is Eq. (4). Eq. (8) arises from a similar reasoning, achieved by expressing the product of exponentials as a single exponential of sums. Although the series $\mathbb{E}[T; n]$ converges for $f > 0$, its closed-form expression is not known. If we rescale time $\tilde{T} \equiv fT$, $\tilde{t} \equiv ft$, and define $x \equiv f/v$, we may rewrite Eq. (S12) in a more compact, non-dimensional form

$$\mathbb{E}[\tilde{T}; n] \equiv \sum_{k=0}^{\lceil (n-3)/2 \rceil} \frac{e^{-xk^2} - e^{-x(k+1)^2}}{2k+1} + \frac{e^{-x(\lceil (n-1)/2 \rceil)^2}}{n}. \tag{S14}$$

As $n \to \infty$, we have

$$\mathbb{E}[\tilde{T}; \infty] \equiv \lim_{n \to \infty} \mathbb{E}[\tilde{T}; n] = \sum_{k=0}^{\infty} \frac{e^{-xk^2} - e^{-x(k+1)^2}}{2k+1} = \sum_{k \in \mathbb{Z}} \frac{e^{-xk^2}}{1 - 4k^2}. \tag{S15}$$

A few interesting observations can be made regarding the upper bounds of this limit.

##### 1.2. On Dawson function estimates

The series $g(x) \equiv \mathbb{E}[\tilde{T}; \infty]$ is related to the family of theta functions (Tyurin, 2002), allowing us to express it in terms of

$$\vartheta(x) = \sum_{k \in \mathbb{Z}} e^{-\pi(xk)^2} \tag{S16}$$

which satisfies $\vartheta(1/x) = x\vartheta(x)$. From Eq. (S15), $g$ satisfies

$$g(x) + 4g'(x) = \sum_{k \in \mathbb{Z}} e^{-xk^2} = \vartheta(\sqrt{x/\pi}), \tag{S17}$$

and thus

$$g(x) = e^{-x/4} \int_0^{x/4} e^y \, \vartheta(2\sqrt{y/\pi}) \, dy. \tag{S18}$$

In particular, for small $x$ we have

$$g(x) = \sqrt{\pi} D_+(\sqrt{x}/2) + O(xe^{-\pi^2/x}) \tag{S19}$$

where

$$D_+(z) = e^{-z^2} \int_0^z e^{t^2} \, dt = \frac{1}{2} \sum_{n=0}^{\infty} \frac{(-1)^n n!}{(2n+1)!} (2z)^{2n+1} \tag{S20}$$

is the Dawson function (Temme, 2010). A less accurate estimate is then $g(x) = \sqrt{\pi x}/2 + O(x^{3/2})$. Various upper bounds may also be obtained this way. Reverting the change of variables, we get $\mathbb{E}[T; \infty] \simeq \frac{1}{2}\sqrt{\frac{\pi}{fv}}$, as in Eq. (5).

### Supplementary Note 2: Computational methods and data

#### 2.1. Beacon Calculus model

As discussed in Boemo et al. (2020), a simplistic model of DNA replication using `bcs` consists of three core process definitions: replication origins (`ORI`), left-moving forks (`FL`), and right-moving forks (`FR`). The origins are positioned along the chromosome of length `L`. Each of these three processes possesses a unique parameter, denoted as `i`, which is assumed to be a specific position on the chromosome between 1 and `L`. In addition to this, the origins have one more parameter: a replication initiation rate, known as `fire`, or $f$ in our model (generalised models may also include licensing probabilities). To monitor which positions on the chromosome have already undergone replication, the model uses markers called beacons. Upon the replication of position `i` by a fork, a beacon is dispatched on the `chr` channel with parameter `i`.

The following is an example of the `bcs` script with 10 replication origins equally spaced over 100 sites

```
// DNA Replication

// Variables
// Chromosome length
L = 100;
// Fast rate
fast = 100000;
// Fork velocity
v = 1.4;

// Process definitions
ORI[i,fire] = {~chr?[i],fire}.(FL[i]||FR[i]);

FR[i] = {chr![i],fast}.[i < L] -> {~chr?[i+1],v}.FR[i+1];
FL[i] = {chr![i],fast}.[i > 0] -> {~chr?[i-1],v}.FL[i-1];

// Process initiation
ORI[1,0.06048832790213383] || ORI[12,0.002045183033099289]
 || ORI[23,0.0012753405213046796] || ORI[34,0.0011945930278953077]
 || ORI[45,0.001035526093646997] || ORI[56,0.0011165358858784408]
 || ORI[67,0.002560893635329413] || ORI[78,0.003411336829553979]
 || ORI[89,0.0022730688407988954] || ORI[100,0.0038028859830789045];

// End
```

A periodic version of DNA replication can be achieved by changing both `FR` and `FL` process definitions to

```
FR[i] = {chr![i],fast}.(([i<L] -> {~chr?[i+1],v}.FR[i+1]) || ([i==L] -> {~chr?[0],v}.FR[0]));
FL[i] = {chr![i],fast}.(([i>0] -> {~chr?[i-1],v}.FL[i-1]) || ([i==0] -> {~chr?[L],v}.FL[L]));
```

##### 2.2. Fitting algorithm

The following code presents the main fitting function, `fitfunction`, used in the fitting algorithm described in previous sections. It provides an efficient way of computing Eq. (8) to mimic `bcs` simulations for non-uniform firing rates. `fitfunction` accepts four arguments: `list` (a data vector with the RT profile of the entire genome), `v0` (average fork speed, usually set to $1.4$ kb/min), and `st0` (parameter $R$, as discussed before). The first guess `x00` is then constructed based on `list`, by Eq. (6). We use an adapted version of `np.roll()`. Data was processed via the Python extension pyBigWig (Ryan et al., 2021). See https://github.com/fberkemeier/DNA_replication_model.git for further details.

```
# Import dependencies
import cProfile
import math
from time import monotonic
from typing import Any
import numpy as np

# Main function
def fitfunction(list, v0, st0):

        timel = list
        v = v0
        st = st0
        exp_v = np.exp(-1/v)
        x00 = np.array([(math.pi/(4*v))*i**(-2) for i in timel])

        # VECTORIZED APPROACH

        def  fast_roll_add(dst, src, shift):
            dst[shift:] += src[:-shift]
            dst[:shift] += src[-shift:]

        def  fp(x, L, v):
            n = len(x)
            y = np.zeros(n)
            last_exp_2_raw = np.zeros(n)
            last_exp_2 = np.ones(n)
            unitary = x.copy()
            for k in range(L+1):
                if k != 0:
                    fast_roll_add(unitary, x, k)
                    fast_roll_add(unitary, x, -k)
                exp_1_raw = last_exp_2_raw
                exp_1 = last_exp_2
                exp_2_raw = exp_1_raw + unitary / v
                exp_2 = np.exp(-exp_2_raw)

                # Compute the weighted sum for each j and add to the total
                y += (exp_1 - exp_2) / unitary

                last_exp_2_raw = exp_2_raw
                last_exp_2 = exp_2
            return y

        def  fitf(time, lst, x0, j):
                return x0[j] * (lst[j]/time[j])**(2)

        def  cfit(time, lst, x0):
            result = np.empty_like(x0)
            for j in range(len(x0)):
                if fitf(time, lst, x0, j) < 10**(-20):
                    result[j] = 10**(-20)
                elif abs(time[j] - lst[j]) < .5:
                    result[j] = x0[j]
                else:
                    result[j] = fitf(time, lst, x0, j)
            return result

        xs = x00
        my_list = [':.20f'.format(i) for i in xs]

        return my_list
```

##### 2.3. Data mappability

Repli-seq data often face mappability issues, particularly in regions with repetitive sequences or low complexity, where short DNA reads cannot be accurately mapped (Hansen et al., 2010; Zhao et al., 2020).

Based on data from Hansen et al. (2010), these regions of low or problematic mappability account for approximately 20% of the whole genome and around 25% of high-error regions (defined as those with errors exceeding $10^2$ min), highlighting their relevance in areas prone to replication timing errors. The mean size of these gaps is approximately 42.37 kb (Figure S1). On average, we observed a phi coefficient of 0.21 when comparing high-error regions and problematic loci, indicating a weak positive correlation between the two. This coefficient, derived from a contingency table, suggests that while there is some overlap between high-error and masked regions, the correlation is not strong. Despite this overlap, mappability issues do not significantly affect overall replication timing analyses, as the majority of high-error regions occur in well-mapped genomic areas, ensuring the reliability of the data.

Given the low phi coefficient, we do not exclude these data from our analysis, since the presence of low mappability regions does not appear to be a major factor influencing replication timing errors, allowing us to retain these data in our analysis without compromising its validity.

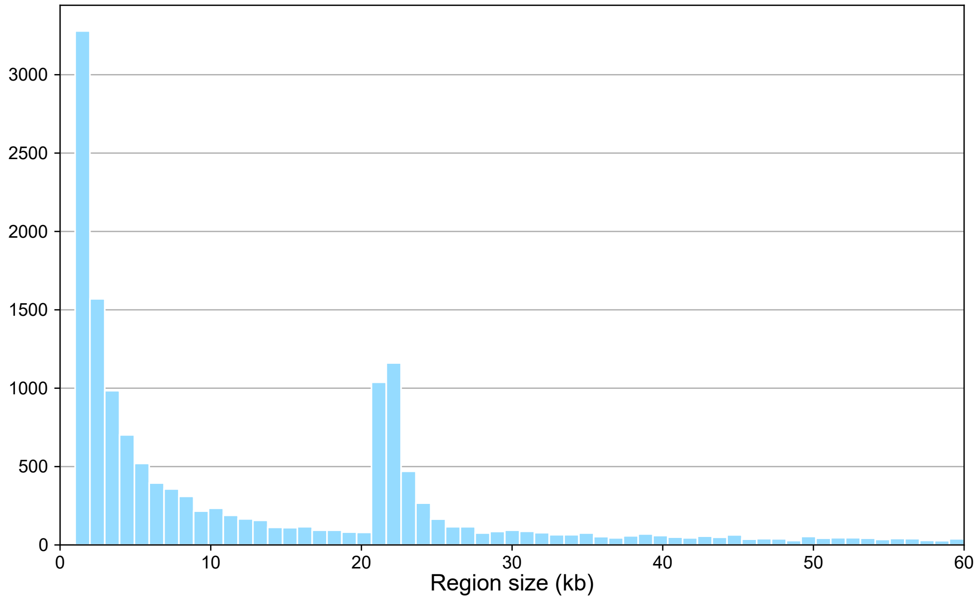


**Figure S1. Distribution of problematic mappability region sizes.**
Histogram showing the distribution of region sizes with low or problematic mappability (in kilobases) across the genome. These regions are excluded from replication timing analyses due to difficulties in accurately mapping sequencing reads. The majority of these regions are small, with peaks around 1-5 kb and another noticeable peak around 20 kb. The mean size of these regions is approximately 42.37 kb.

##### 2.4. Data correlations

Here, we present a comparison of different statistical tests applied to the datasets discussed in the main text. This analysis evaluates the relationships between replication timing error, firing rates, and transcriptional or chromatin features, providing insights into the suitability and results of Pearson, Spearman rank, and Kendall's tau tests for these data.

Pearson, Spearman rank, and Kendall's tau offer distinct advantages based on the nature of the data and relationships analyzed. Pearson is suited for continuous, normally distributed data with linear relationships, while Spearman rank excels with non-linear or ordinal data by capturing monotonic trends through ranked values. Kendall's tau is particularly effective for smaller datasets, using concordant and discordant pairs to measure associations. Given the non-linear and ranked nature of replication metrics, Spearman rank is ideal for our analysis. Figure S2 shows the correlations between replication timing error, firing rates, and transcriptional or chromatin features, demonstrating the relevance of these tests to our data.

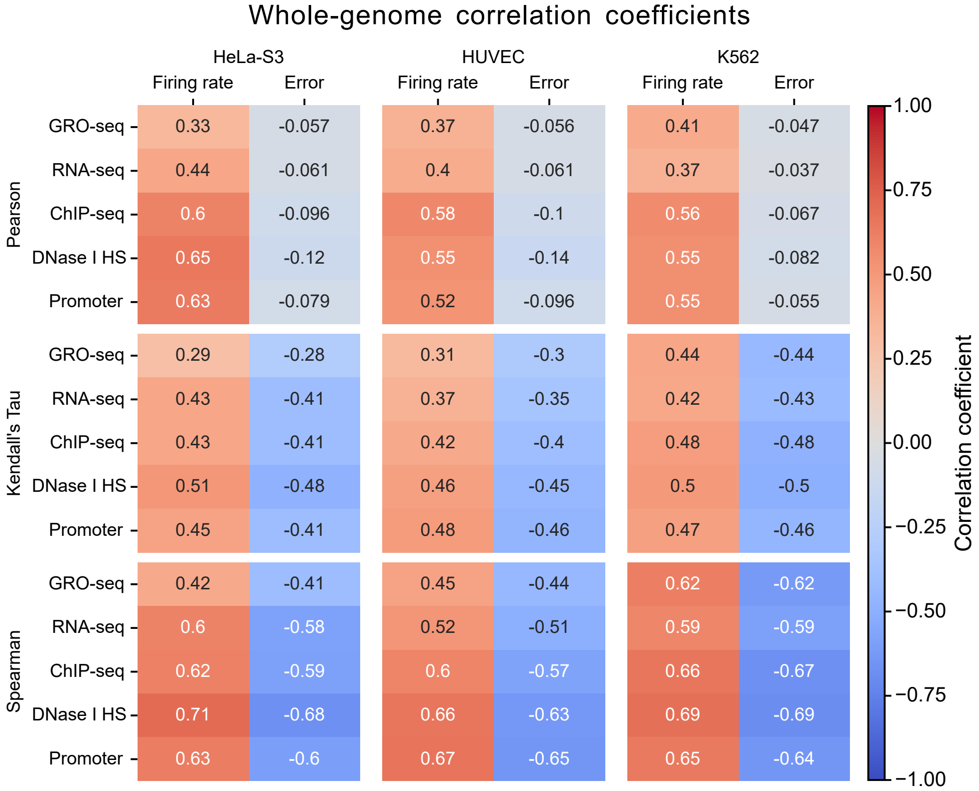


**Figure S2. Correlations between replication, transcription and chromatin data.**
Heatmap displaying the Spearman, Kendall's Tau, and Pearson correlation coefficients between origin firing rates and fit errors with transcriptional and chromatin features for HeLa, HUVEC, and K562 cell lines. All tests returned p-value $< 10^{-15}$.
